# Coping with strangers: how familiarity and active interactions shape group coordination in *Corydoras aeneus*

**DOI:** 10.1101/448068

**Authors:** RJ Riley, ER Gillie, RA Johnstone, NJ Boogert, A Manica

## Abstract

Social groups whose members have had sustained prior experience with each other frequently exhibit improved coordination and outperform groups whose members are unfamiliar with one another. The mechanisms by which familiarity assists coordination are not well known. Prior social experience may simply allow individuals to learn the behavioral tendencies of familiar group-mates and coordinate accordingly. In the absence of prior social experience, it would be adaptive for individuals to develop strategies for coping with unfamiliar others to minimize the disadvantage of unfamiliarity. To explore the dynamics of familiarity in shaping group behaviors, we used a highly social catfish, *Corydoras aeneus*, that utilizes a distinctive, observable tactile interactions. Here we describe this tactile interaction behavior, physical “nudges” that are deployed to initiate group movements and maintain contact with group-mates during group movements. We then report the results of two experiments exploring the relationship between nudges and coordination. First, within triplets of two familiar and one unfamiliar individual, we found no individual differences in nudging rate based on familiarity. Despite all individuals interacting at similar rates, however, unfamiliar individuals failed to coordinate as well as their familiar group-mates, and were more frequently absent from group movements. Second, comparing pairs of familiar with pairs of unfamiliar fish, there was no difference in the level of coordination between pairs. Instead, we found that unfamiliar pairs exhibited significantly higher nudging rates, suggesting that unfamiliar pairs could compensate for their unfamiliarity by nudging more frequently. In contrast, familiar individuals coordinated with comparatively little nudging, presumably because they were experienced with each other. Overall, these results suggest that nudges can be used to improve coordination of group activities, but that their usage is reduced in the case of familiar individuals, implying that these potential signals may be costly.

## Introduction

Animals can gain great benefits from living in groups. Through group coordination, individuals can increase the likelihood of evading predators and improve foraging success (Chivers, 1995, Griffiths, 2004). Familiarity, defined as previous experience with a given other individual, has been shown to increase coordination in a variety of taxa, including birds (Senar, 1990) and schooling fishes (Ward, 2003). For example, great tits show increased anti-predator defenses within groups based on familiarity, with previous experience of nest-site neighbors leading to a greater likelihood of a familiar neighbor joining in to defend a conspecific’s nest (Grabowska-Zhang, 2012). In fathead minnows, familiar groups exhibited greater shoal cohesion and more effective anti-predator behaviors (i.e. predator inspection) in the face of predation threats when compared to unfamiliar groups (Chivers, 1995). The same effect was found in familiar groups of juvenile trout, which responded significantly faster than unfamiliar groups to a predator attack and were more successful foragers (Griffiths, 2004).

Given the benefits of grouping with familiar individuals, it is not surprising that individuals tend to associate preferentially with familiar over unfamiliar individuals in a number of species, including cowbirds (Kohn, 2015) and guppies (Griffiths, 1997). For example, female cowbirds preferentially associate with familiar group-mates when put into a group with familiar and unfamiliar conspecifics (Kohn, 2015). In sticklebacks, the preference for familiar group mates is so strong that individuals tend to prefer smaller groups of familiar individuals to large groups of unfamiliar individuals, even though they generally prefer larger groups to smaller ones (Barber, 2000).

Despite familiarity’s many benefits, the mechanisms by which familiarity improves coordination have rarely been investigated. It seems likely that familiar individuals are better informed about each other’s preferences and characteristics, and thus respond more promptly or appropriately to a partner’s actions. However, quantifying such responses is challenging, as it is often unclear whether individuals are actively trying to coordinate activities or not (King, 2009). In this paper, we study movement coordination in a species of fish, *Corydoras aeneus* (the Bronze Cory catfish), which exhibits an unusual behavior during coordinated activities. In this highly social neotropical catfish, individuals often nudge each other. Individuals deploy this nudging behavior during joint movements both when initiating and during joint movements, thus providing an easily scored behavior that might affect coordination.

We first assessed pairs of individuals to investigate how individuals interacted and coordinated their behaviour when they were not given a choice of partner. We also noted for how much time a given individual was at the front of coordinated movement.

For unfamiliar pairs, we assessed the connection between nudging and coordinated movements. We tested whether pairs’ nudging rates were higher during coordinated movements than when partners were close together but not engaged in coordinated movements. We also investigated whether the amount of time an individual spent as the ‘front fish’ in a coordinated movement was related to the rate at which they nudged their partner.

We then examined both familiar and unfamiliar pairs to test how coordination, cohesion, and the use of nudging varied based on familiarity. We predicted that unfamiliar pairs would require a higher nudging rate to achieve the same level of coordination as familiar pairs.

Finally, we observed the movements of triplets of fish composed of two familiar individuals and one unfamiliar one. We tested whether familiar individuals spent more time close to each other (i.e. coordinated their movements better) than unfamiliar individuals. We also measured the rate at which individuals nudge each other during joint movements, to test whether familiarity affected an individual’s use of this behavior. We predicted that individuals would coordinate better with familiar group-mates.

## Methods

### Corydoras aeneus

*Corydoras* catfish is a highly social neotropical genus widely used in the aquarium trade. In captivity, they have lifespans from 10-15 years (Lambourne, 1995), but their life histories in the wild are not fully known. *Corydoras* are generally benthic fish that prefer slow moving, shallow water; they are known for their marked sociality and shoaling behavior (Nijssen in Lambourne, 1995). In the wild, *Corydoras aeneus*, commonly known as Bronze Cory catfish, are social foragers that live in mixed groups of males, females, and juveniles (Nijssen in Sands, 1986). *Corydoras aeneus* has a slight sexual dimorphism, with females being larger and thicker-bodied than males (Kohda, 2002). We have observed that captive-bred individuals exhibit an unusual tactile interaction behavior during coordinated activities. Wild fish were also observed utilizing this behavior in several small streams in the Madre de Dios locality of the Peruvian Amazon (Riley, pers. obs.).

### Social housing husbandry

We obtained Bronze Cory catfish from three local pet shops in Cambridgeshire: Maidenhead Aquatics Cambridge, Pet Paks LTD, and Ely Aquatics and Reptiles. All fish used in both experiments were at least 24 weeks of age, and had been housed in the lab for at least six weeks prior to the start of experiments. We maintained the fish on reverse osmosis (RO) water purified to 15 or less total dissolved solids (TDS) and re-mineralized to 105-110ppm TDS using a commercially prepared RO re-mineralizing mix (Tropic Marin Re-mineral Tropic). The fish lived on a 12:12 light:dark cycle at a temperature of 23 ± 1 °C. Prior to the start of the experiment, we housed the fish in mixed-sex social housing tanks (60cm × 30 cm × 34 cm) of 6-10 fish. The tanks were equipped with four Interpet Mini internal filters and an air stone. We fed the fish daily on a varied diet of alternating Hikari wafers (Hikari brand, USA), Tetra Prima granules (Tetra brand, Germany), and thawed frozen bloodworms (SuperFish, UK). The group composition of social housing tanks was stable for at least six weeks prior to experiment, and unfamiliar fish had not been exposed to each other for at least six months prior to the experiment, if at all. At the conclusion of each experiment, all fish were returned to the social housing tanks.

### Pair study experimental procedure

We investigated the behavior of familiar and unfamiliar pairs of fish; these trials were completed in three batches. We analyzed 27 pairs for a total of 54 individuals. Each batch consisted of 5-6 familiar pairs, and 5-6 unfamiliar pairs. Experimental batches were tested in October 2016, November 2016, and February 2017 to allow new fish to habituate to the laboratory environment. We formed experimental pairs by randomly assigning individuals to ‘familiar’ or ‘unfamiliar’ treatments. Individuals in the ‘familiar’ condition were paired with an individual from their same social housing tank; unfamiliar individuals were paired with an individual from a different social housing tank (i.e. had not been exposed to each other for at least six months, if ever). As in the case of triplets, we paired individuals in both treatments with same-sex partners to avoid courtship interactions, and fish were not fed prior to trials.

Pairs were placed into one of two filming tanks, which had dimensions (45.5 cm × 25 cm × 21cm) and a sand substrate. Each filming tank had two small plastic plants in one corner of the tank to provide cover. During each session, one filming tank contained a familiar pair while the other contained an unfamiliar pair. Pairs were filmed in random order and were assigned to a filming tank randomly. Pairs were not fed prior to filming. We filmed each pair with a Toshiba Camileo x100 video camera for one-hour.

### Triplet study experiment procedure

We investigated the behavior of familiar and unfamiliar individuals in triplets over three weeks in May-June 2017. We analyzed 18 triplets for a total of 54 individuals. Each triplet consisted of two familiar individuals taken from the same social housing tank and an unfamiliar individual taken from a different tank. Triplets were composed of same-sex individuals to avoid courtship interactions, and fish were not fed prior to the trial to encourage exploratory movement of the tank in search of food. We placed each triplet in one of two testing arenas with a thin layer of aquarium sand as substrate (to reflect the Bronze Cory catfish’s natural habitat). Each arena (47cm × 30 cm × 29 cm) was constructed by partitioning a larger aquarium with a fitted opaque plexiglass sheet. We used a GOPRO HERO 3 camera to film each triplet for 30 minutes from above.

### Video scoring

Triplet and pair videos were scored using the same criteria, except where noted. Video scoring commenced at the first joint trip, which we defined to be a directional group movement (in which each fish was within two body lengths of another individual, and all members of the triplet or pair were moving in the same direction) lasting at least five seconds. Videos were scored by Beth Gillie and Riva Riley, and pair videos were scored blind (i.e. without knowing if any given pair were familiar or unfamiliar). On average, 40% of videos (randomly determined) were scored by both scorers to ensure consistency (for details, please see appendix). The remaining videos were scored by one scorer. We scored triplet videos for five minutes and pair videos for 10 minutes following the first joint trip.

We quantified cohesion by estimating the amount of time two individuals spent in proximity to one another, defined as within 7 cm (roughly two body lengths of an average sized fish). We also recorded “nudges” (tactile interactions), which include any time fish touch one another while in motion, while one is at rest, or when fish are resting close to one another and one individual starts to move in a way that causes it to brush against its partner. For each interaction, we identified the actor and the recipient of the interaction. Since interactions can only occur when fish are in proximity to one other, we focus on the rate of tactile interactions delivered by one fish to another whilst in proximity (i.e. within 7 cm). We define this ‘nudging rate’ as the number of nudges initiated by an individual divided by the number of seconds the pair spend in proximity (i.e. time together).

For pairs, it was possible to classify which individual was in front during coordinated movement (defined as both fish moving in the same direction with one individual at least one half of a body length in front of the other for at least three seconds). In unfamiliar pairs, we investigated whether coordinated movements were associated with a higher nudging rate when compared to individuals close to one another but not engaged in coordinated movements. We also tested whether individuals that initiate more nudges tend to spend more time in front during coordinated movements.

We compared the amount of time in proximity between pairs of familiar and unfamiliar fish (proportion of time together, arcsin(sqrt) transformed), and their rates of tactile interactions (nudges) using two-sample t-tests. As some values seemed to deviate from the assumption of normality, we confirmed our results also by running a Wilcoxon test. Finally, we tested for a link between proportion of time spent in front (arcsin(sqrt) transformed) and nudging rate with a linear model with an interaction between familiarity and initiation rate, with pair ID as a blocking factor. Similarly, for triplets we used an ANOVA test for differences between familiar and unfamiliar fish in the proportion of total time they spent in proximity to one another (we defined proportion of time together as (time together / total time), which was arcsin(sqrt) transformed), and the rates of initiating and receiving tactile interactions, using triplet ID as a blocking factor.

Data were analyzed in R (R version 3.2.2).

## Results

### Nudges and coordination in unfamiliar pairs

Nudging was associated with coordinated movements (when one fish was in front of the other) in unfamiliar pairs: nudging rates during coordinated movements were significantly higher than rates when fish were in proximity with one other but were not coordinating their movements (Figure 2, Paired Wilcoxon signed rank test, df=25, V = 66, p <0.001). There was no association between the amount of time an individual spent in front and its rate of initiating nudges (Pearson’s correlation, t = −0.08, df = 22, p-value = 0.939).

**Figure 1:**
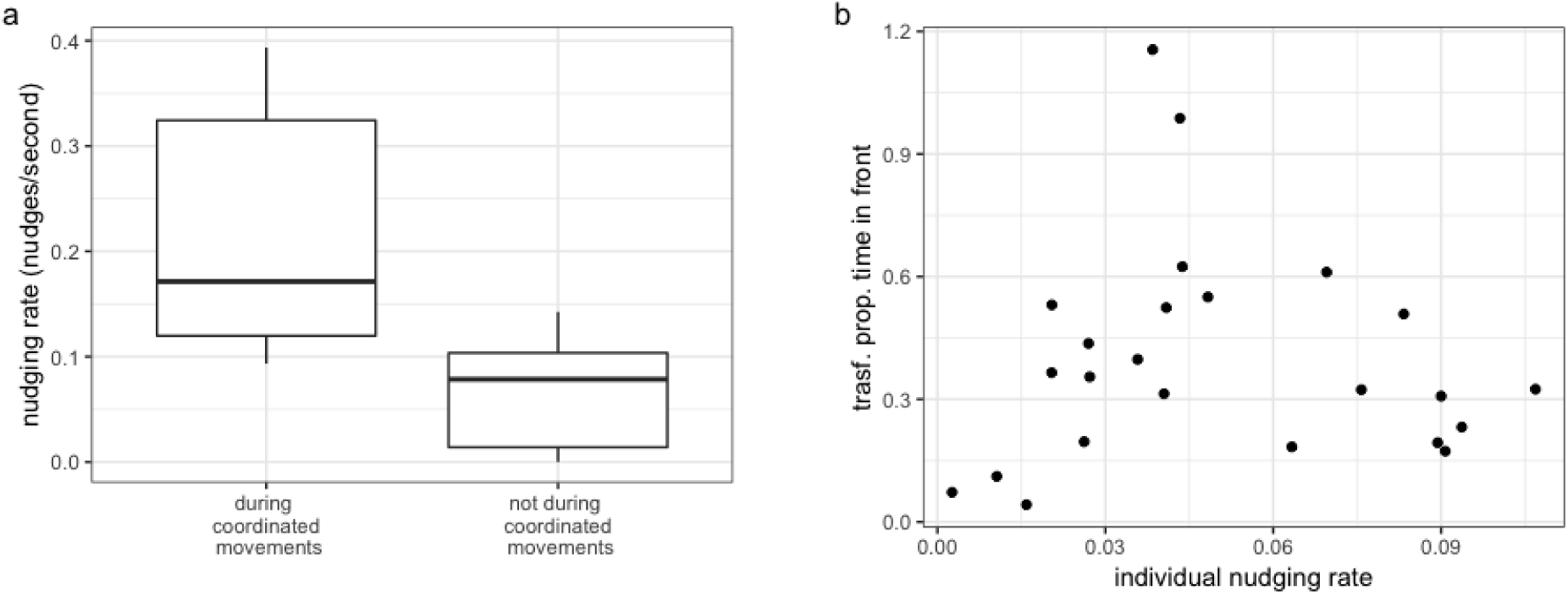
a) nudging rates during and outside of coordinated movements in unfamiliar pairs; b) proportion of time in front (arcsin(sqrt) transformed) vs rate of nudges initiated by the individual in front in unfamiliar pairs.

**Figure 2:**
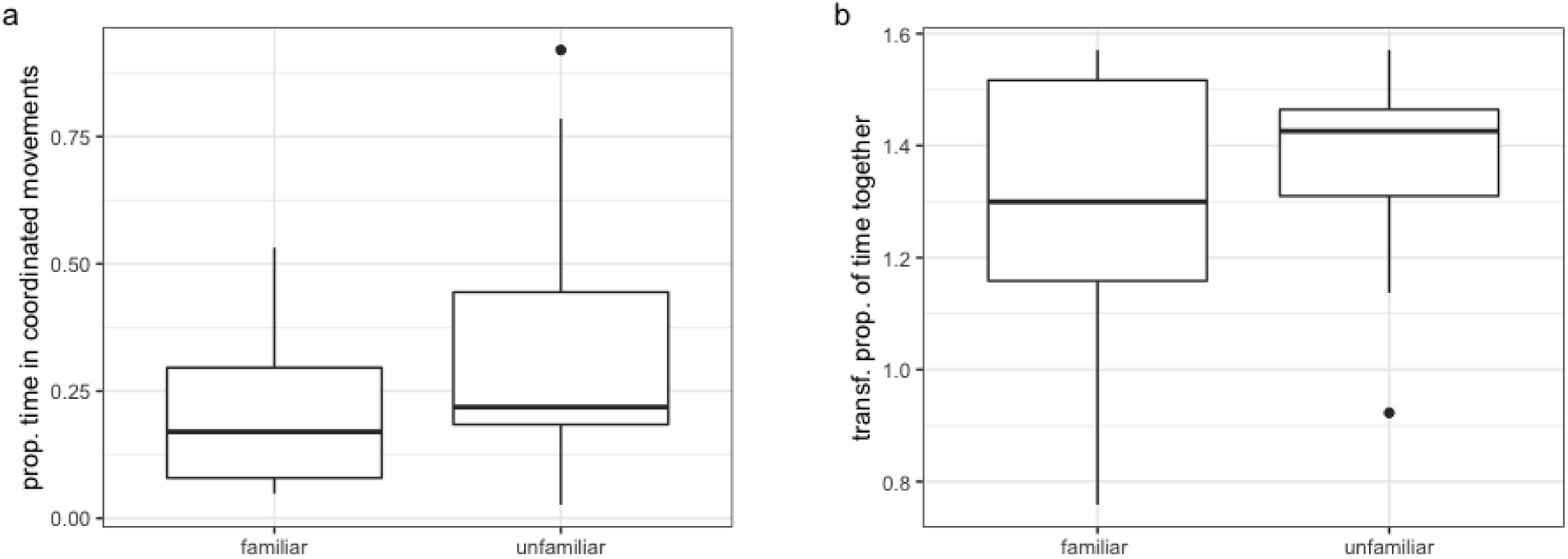
a) proportion of time in coordinated movements in familiar and unfamiliar groups and b) arcsin(sqrt) transformed proportion of time together in familiar and unfamiliar groups

### Comparing coordination and cohesion between familiar and unfamiliar pairs

Familiarity had no effect on the level of coordination or cohesion in pairs of fish. Familiar and unfamiliar pairs spent the same proportion of time engaged in coordinated movements (two sample t-test, t_25_= 1.3, p=0.367, figure 2a) and the same proportion of time in proximity to one another as unfamiliar pairs (two sample t-test, t_25_= −0.93, p= 0.36, fig 2b).

While patterns of coordination and nudging were similar in familiar and unfamiliar pairs, there was a significant difference in familiar and unfamiliar pairs in the rate of nudging, with individuals in unfamiliar pairs nudging each other more frequently than in familiar pairs (two-sample t-test: *t*_25_ = −2.18, p = 0.039, Fig. 3).

**Figure 3:**
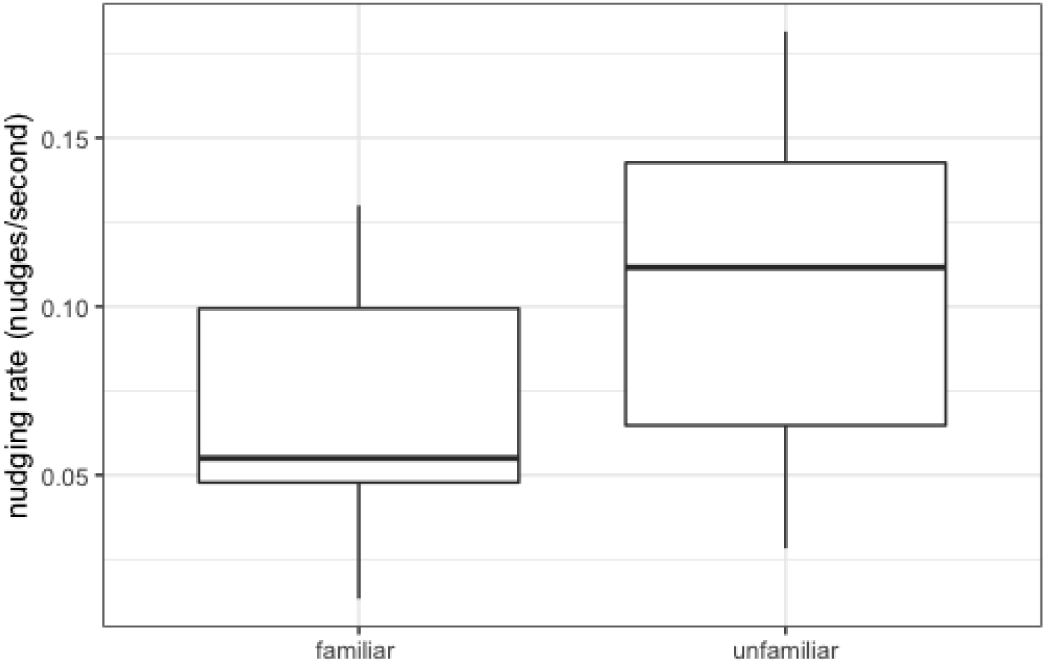
nudging rate in familiar and unfamiliar groups

Figure 3: nudging rate in pairs. Lines inside boxplots indicate medians, boxplot boundaries indicate interquartile range, whiskers indicate +/- 1.5IQR, and points beyond the whiskers are indicated as outliers

### Triplets

Within triplets, familiarity was associated with higher levels of coordination: the two familiar fish spent a higher proportion of time in proximity to one another than they did near the unfamiliar fish (ANOVA, F_1,18_=14.1, P=0.0006, Fig. 4a; removing the outlier does not impact the result). However, when fish were in proximity to one another, there was no effect of familiarity on the rate at which they nudged each other (ANOVA, F_2,18_=2.08, p= 0.131, Fig. 4b).

**Figure 4:**
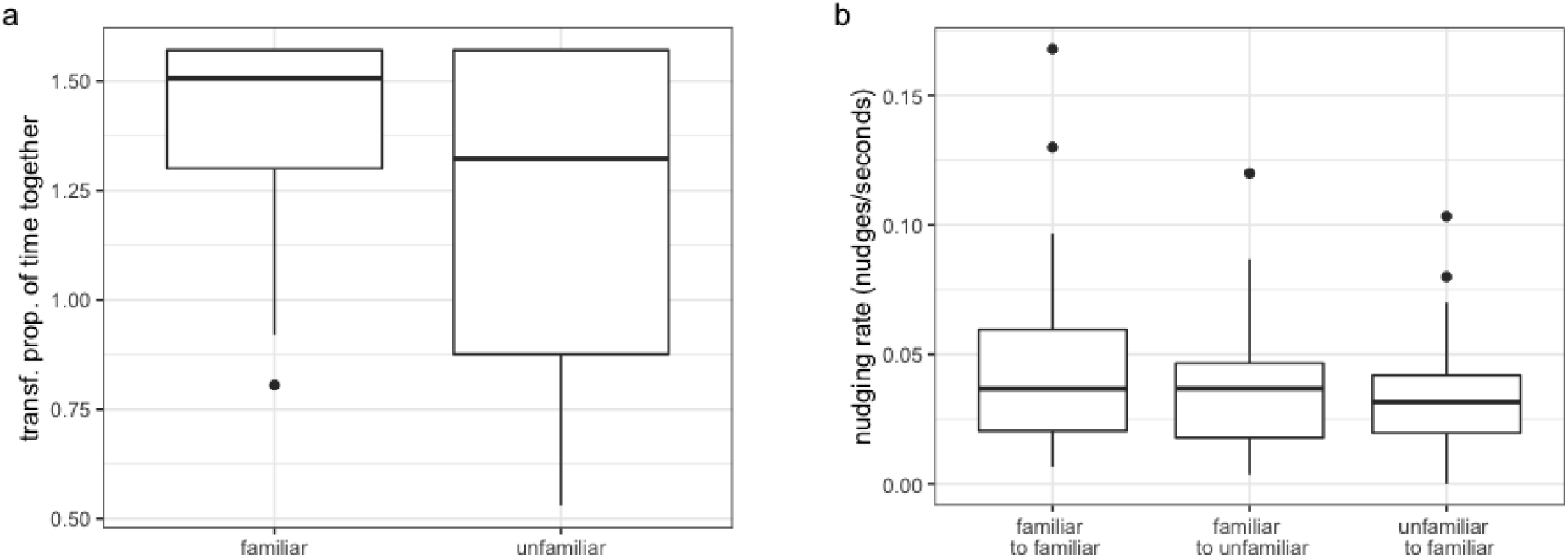
a) arcsin(sqrt) transformed time together for familiar and unfamiliar individuals in triplets; b) pairwise comparisons of nudging rates within triplets. Lines inside boxplots indicate medians, boxplot boundaries indicate interquartile range, whiskers indicate +/- 1.5IQR, and points beyond the whiskers are indicated.

## Discussion

The presence of easily identifiable nudges in Bronze Cory catfish allows us to quantify the link between coordination and interactions among group members (even though we note that nudges are only one method of interacting). Specifically, nudging rates are significantly higher during coordinated movements than when pairs are not engaged in a coordinated movement. The fact that an individual’s nudge initiation rates are not correlated with the individual’s time in front suggests that nudging is not used exclusively by leaders for recruitment, but by both the ‘front fish’ and the ‘back fish’ to maintain coordination during joint movements. This suggests that nudging is a behavior that Bronze Cory catfish individuals utilize while coordinating with others during exploration of a new area, as the pairs and triplets in this study were obliged to do. This is consistent with other systems in which communication behaviors regulate group coordination, as in mouse lemurs, which use olfactory signals to regulate inter-group spatial coordination and acoustic signals to regulate intra-group cohesion and coordination (Braune, 2005). Similar examples exist in birds, for example, in green woodhoopoes, vocalizations are used to maintain group cohesion while moving to new territory (Radford, 2004). The use of tactile interactions by Bronze Cory catfish introduces interactions that utilize a different sensory modality than these examples, and which are strongly associated with coordinated movements directed by a front fish. Consequently, we used nudging as one metric to assess how familiarity affects coordination in triplets and pairs of Bronze Cory catfish.

We found that familiarity affected individuals differently based on group size. Individuals in pairs do not have a choice of group-mates, and without this choice, familiar and unfamiliar pairs spent similar amounts of time together and exhibited similar patterns of nudging and coordination. However, it appears that unfamiliar pairs had to engage in a significantly higher nudging rate in order to achieve the same degree of coordination as familiar pairs. In triplets, however, our results were consistent with the established literature on the effect of familiarity on group coordination. Individuals in a triplet with one familiar group-mate and one unfamiliar group-mate spent more time in proximity with their familiar group-mate, although they nudged both group-mates at similar rates. The effect of cohesion on familiarity is in line with our expectations, and similar results obtained in a number of other species (Chivers, 1995, Griffiths, 1997). This difference between the two experiments reveals that Bronze Cory catfish have the ability to coordinate effectively with unfamiliar individuals, but their failure to do so when given a choice suggests that coordination with unfamiliar individuals is likely to carry some cost (Bronze Cory catfish are highly social, and rarely forage in isolation). Thus, it seems that individuals are willing to pay such a cost only when they have to.

The higher nudging rates in unfamiliar compared to familiar pairs suggest that these “nudges” might play a role in aiding coordination between individuals without prior experience (and therefore without social information) about one another. The finding that increased use of nudges occurred only when individuals were forced to coordinate with an unfamiliar partner suggests that there might be some cost associated with this behavior. Cory catfish rely on camouflage to avoid predators, and freeze when threatened. Thus, it seems likely that individuals might avoid excessive use of nudges, as this is likely to make them more conspicuous to predators. It is somewhat surprising that unfamiliar fish in the triplet experiment did not attempt to increase their interaction rates to help them coordinate with the other two shoal mates; however, it might be the case that, since familiar fish preferentially spent time together, the unfamiliar fish had limited opportunities to interact extensively with them and preferred instead to rely on being still and inconspicuous. Thus, an increased level of nudging was only present in the pair setting, in the absence of a more desirable, and possibly more receptive, partner.

Because unfamiliar pairs coordinate just as effectively as familiar pairs, but require more nudges, while familiar individuals in groups coordinate more effectively with familiar group-mates despite having similar nudging rates with familiar and unfamiliar group-mates, it seems that familiarity may reduce the level of interaction necessary to achieve effective coordination. This suggests that familiar individuals in groups may be able to achieve greater levels of coordination because they have had more chances to interact with one another previously and can respond to one another more effectively. Individuals can then initiate and respond to nudges more effectively based on the previous interactions they have had with their familiar group-mates.

Given the fact that the pairs in our study could achieve the same level of coordination via either familiarity or increased nudging, it is interesting to consider the effect of previous interactions on an individual’s response to conspecifics. Evidence from guppies and deer mice suggests that familiar groups are capable of social learning at a faster rate than unfamiliar ones (Swaney, 2001, Kavaliers, 2005), which could be due to increased cohesion and individuals’ (both demonstrator and follower) greater receptiveness to familiar group-mates. In addition, familiarity reduces aggression in many species, with an individual less likely to display aggressive behaviors toward an individual with which it has prior experience (Ward, 2003, Utne-Palm, 2003, Johnson, 2010). It would be intriguing to investigate how familiarity leads to such outcomes. Familiarity may lead to greater sensitivity to others, which in turn increases social learning potential and reduces the risks of competitive interactions.

Finally, our study highlights the wider importance of exploring active responses to unfamiliarity. There is substantial literature exploring the negative effects of unfamiliarity on groups, but many animals can use tactics to ameliorate these effects. Our results suggest that species that can actively coordinate with potential group-mates will selectively employ such tactics to obviate the disadvantages of unfamiliarity-individuals will do so only when the costs of poor coordination associated with unfamiliarity outweigh the costs of active mediation to coordinate with unfamiliar conspecifics. The costliness of unfamiliarity (likewise, the benefits of familiarity), as well as the necessity for animals to coordinate with others, are a potential selection pressure for the evolution of efficient communication systems.

## Ethical statement

This study was approved via a non-regulated use of animals in scientific procedures application (consistent with UK’s animal welfare legislation ASPA) through the University of Cambridge. It was approved through the University of Cambridge’s Ethical Review Process; it was approved and presented by the Named Veterinary Surgeon and the Named Animal Care and Welfare Officer (NACWO) for the Zoology department.

## Acknowledgements

This study was funded by a Herchel Smith Postgraduate Fellowship.

